# Can we trace the social affiliation of rooks (*Corvus frugilegus*) through their vocal signature?

**DOI:** 10.1101/2024.01.30.577907

**Authors:** Killian Martin, Francesa M. Cornero, Emily Danby, Virgile Daunay, Louise Nervet, Nicola S. Clayton, Nicolas Obin, Valérie Dufour

## Abstract

Inter-individual recognition is crucial for stable social relationships and it is frequently mediated through vocal signatures. In socially complex species, recognition may additionally require additional levels corresponding to other layers of social organisation such as the pair, family, social group or colony. Additional vocal signatures may encode these different levels of social organisations for recognition. We investigated this hypothesis in the calls of the rook (*Corvus frugilegus*), a highly social corvid. Rooks form large breeding colonies where multiple pairs nest in clusters. We recorded the calls of five colonies located in France and in Great Britain, including both wild and captive colonies. To exclude variations due to different call types, we focused on the loud nest call produced exclusively by nesting females during the breeding season. We compared the acoustic distance of calls from each individual and between individuals at various levels of nest proximity, i.e. from the same nest cluster, from different nest clusters, from colonies within the same country, and from colonies in different countries. The only vocal signatures we found were at the individual level, but not at the nest cluster or colony level. This suggests a lack of vocal convergence in this species, at least for the nest call, which may be important for pair recognition in large colonies. Further studies should now evaluate if types of calls other than the nest call better carry vocal signatures as markers of different layers of sociality in this species, or if vocal divergence is a more general vocal phenomenon. In that case, applying new methods of monitoring vocal signatures in wild individuals should help understand the cognitive, social and environmental mechanisms underlying this vocal singularisation.

**Significance statement:** Inter-individual recognition is crucial for social relationships in animals, and is often mediated by individual-specific acoustic characteristics in vocalisations, called a vocal signature. High levels of social organisations, such as a social group of familiar conspecifics or a breeding colony, may likewise be signalled by vocal signatures shared by multiple individuals. We used machine-learning techniques to investigate vocal signatures at multiple social levels in the nest call of brooding female rooks, a corvid species that breeds colonially but lives year-round in social groups. We find evidence of a strong individual vocal signature, but no common vocal signature even in females that nest close together, or in the same colony. A strong individual vocal signature may be a potent tool to monitor populations in this species with minimal disturbance and minimal material, especially as corvids are frequently targeted by human-fauna conflicts in continental Europe.

## 2. Introduction

The ability to recognise different individuals across multiple interactions is crucial to the formation of inter-individual relationships (Bradbury and Vehrencamp, 1998). In many vocally communicative species, conspecifics can be recognised through a vocal signature (Konishi, 1985; Marler, 2004) that corresponds to a set of distinctive vocal characteristics. The abundance of literature on vocal signatures attests to the ubiquity of this phenomenon (e.g. Beecher, 1989; Benti et al., 2019; Bradbury and Vehrencamp, 1998; Elie and Theunissen, 2018; Kershenbaum et al., 2014; Linhart et al., 2019; Průchová et al., 2017; Terry et al., 2005). Vocal signatures are especially important when visual information is insufficient, such as when individuals cannot see one another (Candiotti et al., 2012), live in dense social groups (Catchpole and Slater, 2008), or are morphologically similar (Curé et al., 2009).

Vocal signatures can also arise at levels beyond the individual, such that some vocal characteristics may be common to entire social groups. This acoustic similarity between individuals can arise from several mechanisms. The acoustic adaptation hypothesis (Hansen, 1979) posits that habitat characteristics can promote similar vocal characteristics between individuals. For instance, environments with many obstacles (e.g. trees in forests) can lead individuals to adopt lower-pitched vocalisations to compensate for sound attenuation, while higher-pitched vocalisations can be used in urban environments with many low-pitch noises (Deoniziak and Osiejuk, 2019; Slabbekoorn and Peet, 2003). Genetic proximity can also explain acoustic similarity, as relatives may have closer morphologies than non-relatives. Finally, vocal learning between social partners is also a frequent source of acoustic similarity. Vocal learning can take multiple forms, such as vocal sharing through mimicry (Baker et al., 2000; Kremers et al., 2012), or vocal convergence through the modification of vocalisations to attain a common acoustic structure (Candiotti et al., 2012; Monteiro et al., 2021; Sewall, 2009). When vocalisations are shared throughout a sympatric population but follow macro- or micro-geographic variations, they give rise to dialects (Catchpole and Slater, 2008; Marler and Tamura, 1962; McGregor, 1980; Mundinger, 1982; Searcy, 1992; Wright and Dahlin, 2017). Dialects and other group-level vocal signatures may thus provide information on the geographical, but also social origin of individuals, as well as their familiarity with the conspecifics with whom they interact (Hausberger et al., 2008; Salinas-Melgoza and Wright, 2012).

In some social species, individuals can have different types of relationship according to their social partners, e.g. mating pair, family, foraging partners, social group or colony. Furthermore, the social organisation defined by these relationships may change over time; for instance, fission-fusion systems are defined by stable social groups of several individuals that merge and split for various activities such as breeding or migration (Amici et al., 2008; Archie et al., 2006; Bradbury and Balsby, 2016; Bradbury et al., 2001; Connor et al., 2000; Kelley et al., 2011; Willis and Brigham, 2004). This complex social organisation is frequently associated with higher demands on communicative ability, a link known as the social complexity hypothesis (Blumstein and Armitage, 1997; Freeberg et al., 2012; Peckre et al., 2019). In a complex social system, an individual may belong to several layers of social organisation, and thus need to signal not only their individual identity but also the various social layers they are currently involved in (Kondo and Watanabe, 2009; Vignal et al., 2008). Consequently, their vocalisations might contain acoustic signatures for each of these social layers.

Birds, with their diverse social aggregation patterns, are ideal species to study the link between the social system and the existence of several layers of social and individual vocal signatures (Grueter et al., 2020). Amongst bird species, corvids are compelling models, and particularly the rook (*Corvus frugilegus*). Rooks follow a complex social organisation that includes fission-fusion dynamics (Boucherie et al., 2016; Clayton and Emery, 2007; Coombs, 1960; Patterson et al., 1971; Röell and Bossema, 1982) and several more layers of social complexity. Indeed, rooks are monogamous, with mating pairs focusing most of their activities around each other (Boucherie et al., 2016). Social groups composed of several mating pairs live together year-round (Coombs, 1960; Patterson et al., 1971). Multiple social groups frequently congregate for roosting or breeding, forming colonies of up to thousands of individuals (Clayton and Emery, 2007; Marshall and Coombs, 1957). In addition, rooks are a partially migratory species; some populations are mostly sedentary (e.g. in Northern Europe, where only 12% of individuals migrate, Busse, 1969; Swingland, 1977), while other populations can migrate over 1000 km from one year to the next (e.g. in Western Europe, where up to 63% of individuals migrate, Busse, 1969). Rooks can therefore be expected to be highly socially complex, not only due to the number of different relations individuals can form, but also because their social environment is highly variable throughout the year.

Furthermore, multiple pairs of rooks usually nest in clusters on the same trees, and engage in reciprocal nest defence against other rooks from different nest clusters in the same colony (Coombs, 1960). It remains unclear if this is because individuals from the same nest clusters were already familiar with one another, or whether individuals that do not know each other could be nesting within the same cluster. Furthermore, rooks have been hypothesised to reuse the same nest over consecutive years (Coombs, 1960; Goodwin, 1955; Richardson et al., 1979; Dufour and Clayton, personal observations in captive rooks). It is therefore possible that rooks from the same colony might also be familiar with one another, and so we might expect them to need to recognise each other not just at an individual level, but also at the nest cluster or even colony level.

Corvids, including rooks, are also highly vocal songbirds with rich vocal repertoires (see e.g., Bayens, 1981; Chamberlain and Cornwell, 1971; Ha et al., 2003; Heinrich, 1988; Roskaft and Espmark, 1982). Vocal signatures (see e.g. Benti et al., 2019; Laiolo et al., 2000; Mates et al., 2015; Stowell et al., 2016; Yorzinski et al., 2006) and individual vocal recognition by the birds themselves (Hopp et al., 2001; Kondo et al., 2010; Røskaft and Espmark, 1984; Wascher et al., 2012) have been demonstrated in several corvid species. In rooks, different individual males have their own vocal signature in their most common call (Benti et al., 2019), but this call is produced in multiple contexts, including when the birds move. We aimed to study the vocal signature in a call that could be used to monitor wild birds with minimal disturbance. We selected a call exclusively produced by nesting females during the breeding season (hereafter the “nest call”). The nest call is a powerful vocalisation that is produced while females remain in their nest for several weeks each year while brooding the eggs and caring for the offspring, and which allows individual identification.

We investigated vocal signatures in the nest call in several wild and captive colonies of rooks in England and France. We used a data-driven approach to assess whether the nest call carries elements of individual, group and colonial signatures. Given prior evidence (Benti et al., 2019; Martin et al., 2024), we expected to find signatures at the individual level. Additionally, given that captive birds had lived together for several years, we predicted that birds of both captive colonies would exhibit a stronger colonial signature than the wild birds.

## 3. Material and Methods

### 3.1. Study subjects

We recorded the calls of five colonies of rooks: two wild and one captive in Strasbourg (France), and one wild and one captive near Cambridge (UK). The French wild colonies were recorded on the Cronenbourg university campus (hereafter referred to as the ‘Cronenbourg’ colony) and near the public Parc du Contades (hereafter referred to as the ‘Contades’ colony) in Strasbourg, and the British wild colony (hereafter referred to as the ‘Cambridge wild’ colony) was recorded near Madingley, Cambridge. The captive colonies were housed in outdoor aviaries as part of ongoing scientific projects: the Strasbourg captive colony was housed in the Albert Schweitzer public park (hereafter referred to as the ‘Strasbourg captive’ colony), and the Cambridge captive colony at the Sub-department of Animal Behaviour of the University of Cambridge (hereafter referred to as the ‘Cambridge captive’ colony). All the birds in these two colonies had been caught as fledglings from wild colonies around their respective regions. Notably, some individuals from the captive colonies were captured from the same population as the wild colonies, albeit approximately two decades earlier. The French captive colony had also been housed near the Cronenbourg wild colony until 2018 before being relocated to new premises. All captive birds had been housed together in their respective groups for several years at the time of the study.

### 3.2. Recording protocol

All five colonies were recorded in spring 2022, with recording having started in 2020 for a separate project in the case of the Strasbourg captive colony. We recorded the nest calls of female rooks at the nest during the breeding season with Song Meter 4 recorders (Wildlife Acoustics, USA), using either the built-in internal microphones (Contades and Cambridge wild colonies) or external cable-mounted microphones (3 m cables for both captive colonies, and 20 m cables for the Cronenbourg colony). Microphone gains were adjusted empirically to avoid saturation, and recordings were digitised at 48kHz with a 16-bit rate and stored as uncompressed WAV files.

Audio recordings were carried out between mid-March and mid-April, during the breeding season and before the nests were hidden by leaf growth. Sessions were scheduled between 09:00 and 11:00 and between 13:00 and 15:00. Given that vocalisations seemed less frequent at those times compared to others, this reduced the likelihood of overlaps between vocalising females.

Captive females were identified by coloured leg rings and nest position, and were previously habituated to human presence. Wild females were identified only by their nest position. Human observers located at similar heights to the nests (i.e., standing at ground level for the captive colonies, and on nearby rooftops for the Cronenbourg and Contades wild colonies) monitored and annotated which female was calling and when. For the wild Cambridge colony, individual females could not be reliably identified at the time of the recordings due to leaf growth, thus excluding the use of this colony to uncover individual vocal signatures. All monitoring was opportunistic: as soon as a female was heard vocalising and was clearly identified in her nest, each vocalisation was noted using a custom Python script recording the time of the vocalisation as well as the identity of the female. For the French wild colonies, nest positions and nests clusters were mapped, and the sampled nests were noted down on the map as recording sessions progressed in order to reduce the risk of repeatedly sampling the same females (see Supplementary Material for images of the final sampled nests in the ‘Strasbourg wild’ and ‘Contades’ colonies). We defined clusters of nests as nests located at around the same height and in the same tree, up to a maximum distance of approximately 4 m apart. Videos were also made to help clear up possible ambiguities.

Nest calls were then extracted using the Audacity software (v3.1.3). Only calls that did not overlap other sounds were used. This resulted in a total of 2859 calls (Table 1).

**Table 1:** Distribution of nest calls in the study data, per colony. Total: number of calls collected; Mean ± SD: average and standard deviation of calls per individual female; Min-max: minimum and maximum number of calls extracted per individual female; Individuals: number of individual females monitored in the colony.

| Colony | Total | Mean ± SD | Min - Max | Individuals |
| --- | --- | --- | --- | --- |
| Strasbourg captive | 479 | 79.8 ± 61.5 | 9 - 194 | 6 |
| Cronenbourg | 941 | 24.1 ± 20.7 | 6 - 94 | 39 |
| Contades | 283 | 15.7 ± 8.13 | 7 - 35 | 18 |
| Cambridge captive | 223 | 55.8 ± 4.03 | 51 - 60 | 4 |
| Cambridge wild | 933 | - | - | - |

### 3.3. Audio representation

Each call was converted to a spectrogram using a 10 ms Hamming window with 80% overlap, denoised by a dynamic spectral gating algorithm (Sainburg 2020), squared, and finally Mel-scaled down to 80 frequency coefficients to reduce dimensionality and emphasise low frequencies, where most of the energy in rook vocalisations is concentrated (Benti et al., 2019; Røskaft and Espmark, 1982). The Mel-scaling was limited to values above 100Hz to remove low frequency noise. The spectrograms were then log-scaled, limited to a 20 dB dynamic range below the maximum magnitude, and min-max normalised between 0 and 1.

### 3.4. Pairwise acoustic distance between calls

The acoustic distance between calls was estimated using the Dynamic Frequency-Time Warping (DFTW) distance (see Martin et al., 2024 for details). This distance finds the optimal alignment between two spectrograms, first along the time axis, then along the frequency axis, and finally along the time axis again. This allows vocalisations that have similar acoustic structures but are produced at different speeds or with different pitches to still be considered similar. The resulting pairwise distance matrix was then used to assess similarity between individual females, groups and colonies.

### 3.5. UMAP projection

For visualisation purposes, we projected the pairwise DFTW distance matrix down to two dimensions using UMAP (McInnes et al., 2018). UMAP aims to preserve neighbourhoods between points (i.e. points that are originally close together should remain close together in the projection). However, the algorithm does not preserve distance values, so we do not report statistical analysis based on its results. We used UMAP with the default parameters in the umap-learn package (v0.5.3), with the exception of setting the minimum distance between points to 0 (thus allowing points to overlap in the projection if the acoustic distance between the correpsonding calls is low enough).

### 3.6. Statistical analysis

We based analysis on the principle of homophily: if vocal signatures exist, then calls with the same signature should be more similar (and thus, have lower acoustic distances between them) than calls with different signatures. Accordingly, this would mean that individual signatures should give rise to clusters of calls by individual females, group signatures should give rise to clusters of calls by cluster of nests, and that colonial signatures should give rise to clusters of calls by colony. We tested this assertion with two different procedures: first by directly comparing acoustic distances between different levels of nest proximity, and second by verifying that calls from a given individual female (respectively nest cluster, colony) should be closer neighbours to calls from the same individual female (respectively nest cluster, colony) than to calls of other individuals (respectively nest clusters, colonies).

We compared acoustic distances between pairs of calls by level of nest proximity with a GLMM (Gamma family, log link), with the individual female, nest cluster and colony associated to each of the pair of calls as a random factor. The model was fitted in R (v4.1.1, R Core Team, 2021) using the *lme4* package (v1.1-30, Bates et al., 2015). This model was compared to a null model without nest proximity using a LRT test, then pairwise tests were conducted between levels of nest proximity using the *emmeans* package (v1.8.0, Lenth, 2023), applying a Tukey correction for multiple testing. We defined six levels of nest proximity: 1) same individual (between calls from the same individual female), 2) same cluster (between calls from different females but within the same nest cluster), 3) same colony (between calls from individuals in different nest clusters but within the same colony), 4) unknown (specifically for females from the Cambridge wild colony, where individual and nest cluster identities were unknown), 5) same country (between calls from different colonies that were in the same country, i.e. between French colonies and between British colonies), and 6) different countries (between calls from the French and the British colonies). The last two levels were defined because rooks within the same country are more likely to be in contact through migration or movements outside of the breeding season than rooks from different countries in this study. Finally, for the captive colonies, only one nest cluster existed for each due to the aviary conditions, but calls from different females were considered as part of the “same colony” level nest proximity instead of the “same cluster”.

The second analysis used a k-nearest neighbours (kNN) approach based on Thomas et al. (2022). In practice, this approach quantifies the probability that, for a call from a given e.g. colony, its *k* nearest neighbouring calls (i.e. calls at lower acoustic distances) are also from the same colony. This probability was then normalised by the same probability under random chance, equal to the proportion of calls from the same colony in the entire dataset. However, this approach may also include calls from the same individual, which may artificially increase probability values if individual vocal signatures exist. To account for this possibility, in a second step we computed the same probabilities while discarding calls from the same individual when counting the k-nearest neighbouring calls.

## 4. Results

All analyses were based on the principle of homophily. We hypothesised that individual (respectively nest cluster, colonial, country) vocal signatures should result in calls from the same individual (resp. nest cluster, colony, country) being acoustically close together, and further from calls from different individuals (resp. nest clusters, colonies, countries). The first GLMM-based analysis indicated that the nest proximity level (i.e. individual, cluster, colony, country) significantly impacted acoustic distance between the nest calls of female rooks (LRT: *x*^2^(*df* = 5)= 28.5, p *<* 0.0001). The post-hoc pairwise tests between nest proximity levels supported the existence of individual vocal signatures: calls from the same female were significantly closer acoustically than calls from different females, whether these other individuals were from the same nest cluster, from the same colony, from different colonies in the same country or from different colonies in the other country (lines 1, 2, 4, and 5 respectively in Table 2). In contrast, there was little to no support for vocal signatures at the nest cluster, colony or country level: the calls from different individuals were not statistically closer whether they were from the same nest cluster, different nest clusters in the same colony, different colonies in the same country or different countries. Only two cases indicated a statistical difference: calls from individuals in the same colony were acoustically closer than calls from different colonies, and calls from individuals from different colonies in the same country were acoustically more distant than those from individuals in different countries. Nevertheless, in both cases, the associated ratios were very close to 1, indicating that the effects were very weak.

**Table 2:** Pairwise comparisons of acoustic distances between levels of nest proximity. Each line corresponds to one comparison, with the coloured cells indicating the specific levels being compared (with the same colour code as in Fig. 1), the ratio (where the numerator is the level in the left cell, and the denominator is the level in the right cell) and standard error (SE), and the associated z-statistic and p-value after Tukey’s correction for multiple testing. Significant p-values and strong effects (indicated by ratio particularly different from 1) are indicated in bold.

| Contrast |  | Ratio | SE | z | p-value |
| --- | --- | --- | --- | --- | --- |
| Same individual | Same cluster | **0.90** | 0.01 | -7.39 | < **0.001** |
| Same individual | Same colony | **0.89** | 0.01 | -11.10 | < **0.001** |
| Same individual | Unknown (Cambridge wild) | 0.99 | 0.32 | -0.03 | 1 |
| Same individual | Same country | **0.87** | 0.01 | -13.10 | < **0.001** |
| Same individual | Different countries | **0.91** | 0.01 | -7.47 | < **0.001** |
| Same cluster | Same colony | 0.99 | 0.01 | -0.68 | 0.99 |
| Same cluster | Unknown (Cambridge wild) | 1.10 | 0.36 | 0.31 | 1 |
| Same cluster | Same country | 0.97 | 0.01 | -2.29 | 0.20 |
| Same cluster | Different countries | 1.01 | 0.01 | 0.66 | 0.99 |
| Same colony | Unknown (Cambridge wild) | 1.11 | 0.36 | 0.33 | 1 |
| Same colony | Same country | 0.98 | 0.01 | -3.63 | **0.004** |
| Same colony | Different countries | 1.02 | 0.01 | 1.76 | 0.49 |
| Unknown (Cambridge wild) | Same country | 0.88 | 0.28 | -0.39 | 0.99 |
| Unknown (Cambridge wild) | Different countries | 0.91 | 0.30 | -0.28 | 1 |
| Same country | Different countries | 1.04 | 0.01 | 3.70 | **0.003** |

We then used kNN analysis to investigate whether acoustic proximity could still be used to identify colonies by the principle of homophily. We hypothesised that a colonial signature should result in vocalisations from each colony forming clusters at a distance from vocalisations from other colonies. However, the kNN analysis results (Fig. 2) did not support this hypothesis. Calls of each colony were indeed more likely than expected by random chance to be within the nearest neighbours of other calls from their respective colony, but controlling for calls from the same individual almost completely cancelled this colonial effect. Finally, a UMAP projection in 2D (Fig. 3) showed that no clusters emerged that separated different colonies, or different nest clusters, and rather indicated that there was a large amount of overlap between different colonies and different nest clusters.

**Figure 1:**
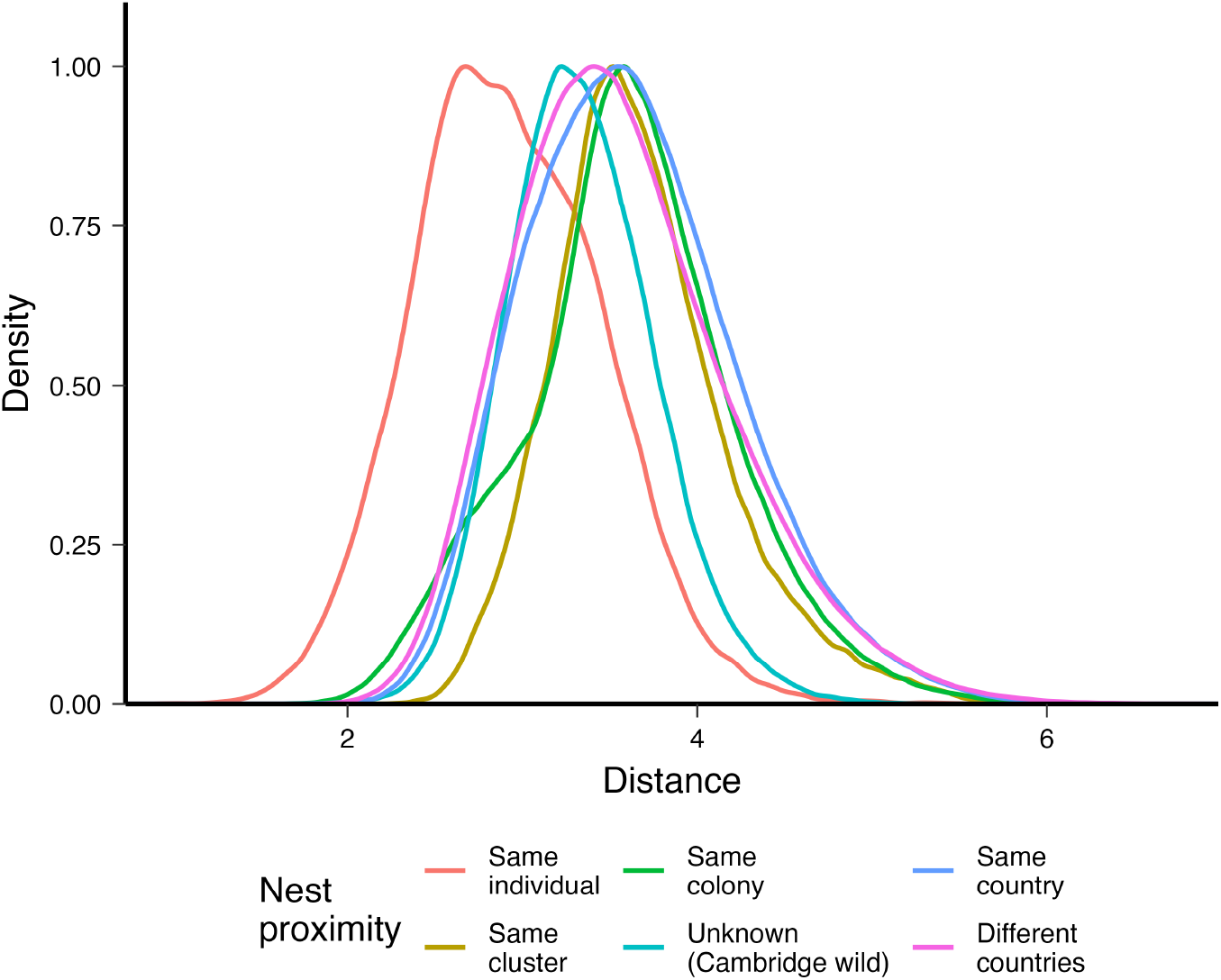
Distribution of the values of acoustic distance between calls for each level of nest proximity. Each line represents the density of the distribution, normalised to the same maximum height. See Table 2 for statistical contrasts between levels.

**Figure 2:**
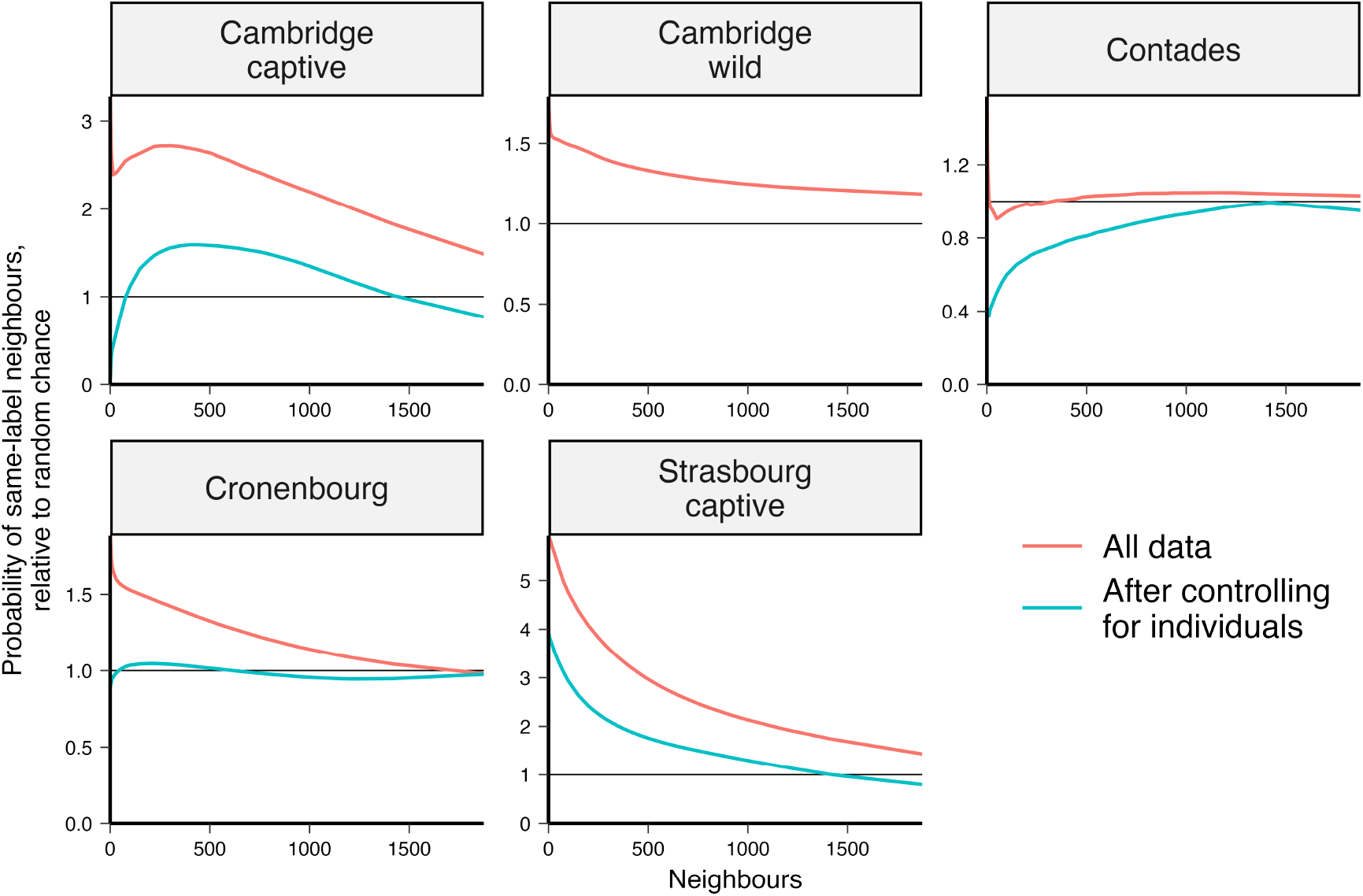
Homophily analysis of calls by colony, before (red lines) and after (blue lines) controlling for individuals. Each line corresponds to the ratio of the proportion of calls from the same colony in the nearest neighbouring calls, over the same proportion expected by random chance. For instance, for the Strasbourg captive colony, the nearest neighbour of calls from this colony is slightly under 6 times more likely than random chance to also be a call from the Strasbourg captive colony, and slightly under 4 times more likely than random chance after correcting for individual signatures. Since individual identity and nest clusters were unknown in the Cambridge wild colony, the corresponding line is not included.

**Figure 3:**
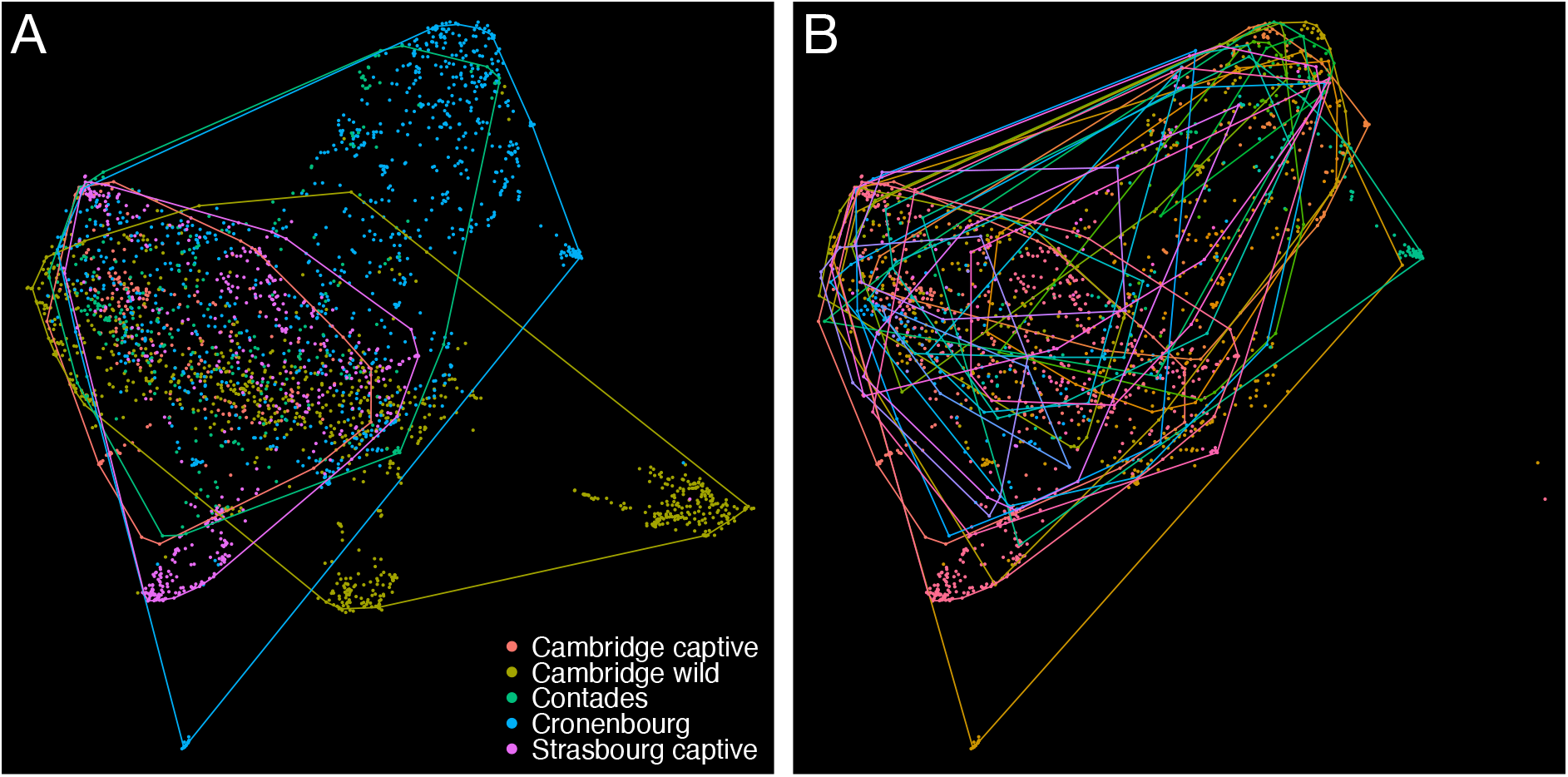
UMAP projection of the DFTW distance matrix. Each point represents one call, coloured by A) colony and B) nest cluster (legend not included to reduce visual clutter). The envelopes were drawn to surround 95% of the vocalisations of each colony and nest cluster, using the same colour code. In B), calls from the Cambridge wild colony were omitted due to the absence of nest cluster information for this colony.

## 5. Discussion

We showed that individual signatures can be detected in the nest calls of female rooks. However, there was little evidence of vocal signatures at the nest cluster or colony levels. Calls from different individuals from the same colony or nest cluster are not more acoustically similar than calls from individuals in different colonies or different nest clusters, and accordingly do not cluster together, as shown by the UMAP projection. An individual signature had already been detected in the most common “caw” calls of male rooks (Benti et al., 2019), although subsequent analysis showed that this could be due to a lack of common call types in the repertoires of male rooks (Martin et al., 2024). Females do however share common call types regardless of their origin, including the nest call. The nest call is a powerful call produced by brooding females when they stay in the nests during the breeding season, and thus depend entirely on their mate to feed them and their offspring for several weeks (Coombs, 1960, 1961). The nest call may therefore be crucial for the pair to recognise and find each other in the large colonies (up to several thousand breeding pairs) that rooks usually form during the breeding season (Coombs, 1960).

The lack of evidence for a colonial vocal signature in rooks was contrary to our hypothesis, given the complex social organisation followed by this species. Rooks live in stable social groups throughout the year (Clayton and Emery, 2007), and nest in clusters which might correspond to these social groups (Coombs, 1960; Goodwin, 1955; Richardson et al., 1979). We therefore expected a phenomenon similar to the dialects often observed in other social songbirds, where individuals that live together converge in their vocal production (Catchpole and Slater, 2008; Mundinger, 1982). The lack of vocal convergence in the present study could have two potential causes. First, individuals within a nest cluster or a colony may not be socially close. Rooks usually disperse back into smaller social groups after breeding, and are frequently exposed to individuals from different groups, including migrating birds (Busse, 1969; Patterson et al., 1971). At the same time, rooks, like several other corvids, remain capable of vocal mimicry during adulthood (Brown and Farabaugh, 1997; Kaplan, 1999). If their vocalisations are influenced by their early social environment, frequent exposure to unknown birds may weaken the influence of the environment at the nest, and thus reduce the occurrence of colonial signatures. Second, vocal convergence may be weakened by the major anthropogenic conflicts involving these species. Approximately four million corvids are killed every year in Europe (Jiguet, 2020), with rooks representing a frequent target of destruction due to the damage caused to crops by their seed-based diet (Feare 1974, 1978; Porter et al 2008). Frequent eradications may cause instability in rook populations by leaving nesting sites open to new incomers at the next breeding season. These new individuals may integrate colonies and decrease population stability over time, possibly preventing vocal convergence within the colony or cluster.

Alternatively, it may be that individuals that live year-round in the same social groups do not vocally converge (at least as far as the nest call is concerned). The strongest argument in favour of a natural lack of convergence is the lack of vocal signature in the two captive colonies despite years of stability in the group composition. We can therefore consider that although population instability may explain part of the lack of colonial vocal signature observed in wild rooks, it is likely not the only explanation. Previous results seem to indicate that vocal singularisation may be more common than vocal sharing in male rooks (Martin et al. 2023). This type of phenomenon, where individual singularisation outweighs a group signature, has been described in the monk parakeets, a species of parrot that also nests in communities (Smith-Vidaurre et al., 2020). However, the lack of vocal sharing or convergence, or even outright vocal avoidance (where individuals actively avoid associating with individuals producing similar vocalisations), has most frequently been described in territorial species (e.g. the chaffinch *Fringilla coelebs*, Slater and Ince, 1982, or the white-eyed vireo *Vireo griseus*, Borror, 1987; Bradley, 1981). It has been suggested that a lack of vocal sharing in these species minimises consanguinity or competition between relatives, but it remains unclear why this phenomenon should arise in a socially breeding species like the rook or the monk parakeet.

Further investigations on vocal signatures are required in rooks, and additional studies should now investigate other call types in this species. New tools for bioacoustic analysis are constantly emerging in the literature, and the latest powerful approaches are based on machine learning (Sainburg and Gentner, 2021). These new approaches, which can take entire spectrograms as input, may permit much more detailed studies of vocal similarity and vocal learning in the wild. Traditional approaches, based on acoustic measures (e.g. Benti et al., 2019; Keenan et al., 2020), are not always appropriate. Rooks, like many other corvids, produce highly noisy and chaotic vocalisations (Fletcher, 2000), which precludes the use of common measures like the fundamental frequency or the harmonic distribution of frequencies (Stowell et al., 2016). Spectrogram-based approaches (see also Sainburg and Gentner, 2021), can bypass this limit and be deployed for wildlife monitoring or to follow population dynamics. In rooks, the nest call does carry markers of female identity, which can be used to follow individual females in long-term studies. Indeed, individual signatures could be used to study social stability in colonial species over time and follow the impact of environmental changes and anthropogenic pressures on bird populations. Moreover, new methods of monitoring wild individuals should facilitate research into the cognitive, social and environment mechanisms underlying this vocal singularisation. More generally, our study promotes non-invasive methods for the investigation of population dynamics, and the link between social complexity and vocal behaviour.

## 6. Ethics

This study was observational only for both wild and captive colonies, and we followed ASAB animal care guidelines for the captives colonies. The rooks in the French colonies and in the wild Cambridge colony required no further approval according to respective country guidelines. The rooks in the Cambridge captive colony were kept in aviary space under the University of Cambridge’s AWERB review and monitoring, and the study was approved as a non-regulated procedure.

## Supporting information

Cartography of wild colonies with sampled nests

## 7 Acknowledgements

This project was supported by a Michelin Fondation donation to V. Dufour (no. 349-PE-3176).

## 8. Data acessibility

The dataset used for this study can be downloaded at https://zenodo.org/records/10580110. The code used for the analysis can be found at https://gitlab.com/kimartin/rook_vocal_signature.

## Notes

https://zenodo.org/records/10580110

https://gitlab.com/kimartin/rook_vocal_signature

### Competing Interest Statement

The authors have declared no competing interest.

## References

Amici, F., Aureli, F., & Call, J. (2008). Fission-Fusion Dynamics, Behavioral Flexibility, and Inhibitory Control in Primates. Current Biology, 18(18), 1415–1419. 10.1016/j.cub.2008.08.020

Archie, E. A., Moss, C. J., & Alberts, S. C. (2006). The ties that bind: Genetic relatedness predicts the fission and fusion of social groups in wild African elephants. Proceedings of the Royal Society B: Biological Sciences, 273(1586), 513–522. 10.1098/rspb.2005.3361

Baker, M. C., Howard, T. M., & Sweet, P. W. (2000). Microgeographic variation and sharing of the gargle vocalization and its component syllables in black-capped chickadee (Aves, Paridae, Poecile atricapillus) populations. Ethology, 106(9), 819–838. 10.1046/j.1439-0310.2000.00602.x

Bates, D., Mächler, M., Bolker, B., & Walker, S. (2015). Fitting linear mixed-effects models using lme4. Journal of Statistical Software, 67(1), 1–48. 10.18637/jss.v067.i01

Bayens, G. (1981). The role of the sexes in territory defence in the Magpie ( Pica pica ). Ardea, 69(1), 69–82. http://ardea.nou.nu/ardea%7B%5C%7Dshow%7B%5C%7Dabstract.php?lang=uk%7B%5C&%7Dnr=763

Beecher, M. D. (1989). Signalling systems for individual recognition: an information theory approach. Animal Behaviour, 38, 248–261.

Benti, B., Curé, C., & Dufour, V. (2019). Individual signature in the most common and context-independent call of the Rook (Corvus frugilegus). The Wilson Journal of Ornithology, 131(2), 373. 10.1676/18-41

Blumstein, D. T., & Armitage, K. B. (1997). Does sociality drive the evolution of communicative complexity? A comparative test with ground-dwelling sciurid alarm calls. American Naturalist, 150(2), 179–200. 10.1086/286062

Borror, D. J. (1987). Song in the white-eyed vireo. Wilson Bull., 99(3), 377–397.

Boucherie, P. H., Mariette, M. M., Bret, C., & Dufour, V. (2016). Bonding beyond the pair in a monogamous bird: Impact on social structure in adult rooks (Corvus frugilegus). Behaviour, 153(8), 897–925. 10.1163/1568539X-00003372

Bradbury, J. W., & Balsby, T. J. (2016). The functions of vocal learning in parrots. Behavioral Ecology and Sociobiology, 70(3), 293–312. 10.1007/s00265-016-2068-4

Bradbury, J. W., Cortopassi, K. A., & CLemmons, J. R. (2001). Geographical variation in the contact calls of orange-fronted parakeets. The Auk, 118(4), 958–972. 10.1093/auk/118.4.958

Bradbury, J. W., & Vehrencamp, S. L. (1998). Principles of Animal Communication (2nd Editio). Sinauer Associates, Inc,.

Bradley, A. (1981). Song variation within a population of white-eyed vireos (Vireo griseus). The Auk, 98(January), 80–87.

Brown, E. D., & Farabaugh, S. M. (1997). What birds with complex social relationships can tell us about vocal learning: Vocal sharing in avian groups. In C. T. Snowdon & M. Hausberger (Eds.), Social influences on vocal development (pp. 98–127). Cambridge University Press. 10.1017/cbo9780511758843.007

Busse, P. (1969). Results of ringing of European Corvidae. Acta Ornithologica, 11, 236–328.

Candiotti, A., Zuberbühler, K., & Lemasson, A. (2012). Convergence and divergence in Diana monkey vocalizations. Biology Letters, 8(3), 382–385. 10.1098/rsbl.2011.1182

Catchpole, C. K., & Slater, P. J. (2008). Bird song: Biological themes and variations, second edition. Cambridge University Press. 10.1017/CBO9780511754791

Chamberlain, D. R., & Cornwell, G. W. (1971). Selected vocalizations of the common crow. The Auk, 88(3), 613–634.

Clayton, N. S., & Emery, N. J. (2007). The social life of corvids. Current Biology, 17(16), 652–656.

Connor, R. C., Wells, R. S., Mann, J., & Read, A. J. (2000). Social Relationships in a Fission-Fusion Society. Cetacean Societies: Field Studies of Dolphins and Whales, 91–126.

Coombs, C. J. F. (1960). Observations on the Rook Corvus Frugilegus in Southwest Cornwall. Ibis, 102(3), 394–419. 10.1111/j.1474-919X.1960.tb08417.x

Coombs, C. J. F. (1961). Rookeries and roosts of the rook and jackdaw in south-west cornwall. Bird Study, 8(1), 32–37. 10.1080/00063656109475986

Curé, C., Aubin, T., & Mathevon, N. (2009). Acoustic convergence and divergence in two sympatric burrowing nocturnal seabirds. Biological Journal of the Linnean Society, 96(1), 115–134. 10.1111/j.1095-8312.2008.01104.x

Deoniziak, K., & Osiejuk, T. S. (2019). Habitat-related differences in song structure and complexity in a songbird with a large repertoire. BMC Ecology, 19(1), 1–11. 10.1186/s12898-019-0255-7

Elie, J. E., & Theunissen, F. E. (2018). Zebra finches identify individuals using vocal signatures unique to each call type. Nature Communications, 9(1). 10.1038/s41467-018-06394-9

Fletcher, N. H. (2000). A class of chaotic bird calls? The Journal of the Acoustical Society of America, 108(2), 821–826. 10.1121/1.429615

Freeberg, T. M., Dunbar, R. I., & Ord, T. J. (2012). Social complexity as a proximate and ultimate factor in communicative complexity. Philosophical Transactions of the Royal Society B: Biological Sciences, 367(1597), 1785–1801. 10.1098/rstb.2011.0213

Goodwin, D. (1955). Some observations on the reproductive behaviour of rooks. British Birds, 48, 97–107.

Grueter, C. C., Qi, X., Zinner, D., Bergman, T., Li, M., Xiang, Z., Zhu, P., Migliano, A. B., Miller, A., Krützen, M., Fischer, J., Rubenstein, D. I., Vidya, T. N., Li, B., Cantor, M., & Swedell, L. (2020). Multilevel Organisation of Animal Sociality. Trends in Ecology and Evolution, 35(9), 834–847. 10.1016/j.tree.2020.05.003

Ha, R. R., Bentzen, P., Marsh, J., & Ha, J. C. (2003). Kinship and association in social foraging northwestern crows (Corvus caurinus). Bird Behavior, 15(2), 65–75. http://www.ingentaconnect.com/content/cog/bb/2003/00000015/00000002/bb063

Hansen, P. (1979). Vocal learning: its role in adapting sound structures to long-distance propagation, and a hypothesis to its evolution. Animal Behaviour, 27(4), 1270–1271.

Hausberger, M., Bigot, E., & Clergeau, P. (2008). Dialect use in large assemblies: A study in European starling Sturnus vulgaris roosts. Journal of Avian Biology, 39(6), 672–682. 10.1111/j.1600-048X.2008.04307.x

Heinrich, B. (1988). Winter foraging at carcasses by three sympatric corvids, with emphasis on recruitment by the raven, Corvus corax. Behavioral Ecology and Sociobiology, 23(3), 141–156. 10.1007/BF00300349

Hopp, S. L., Jablonski, P., & Brown, J. L. (2001). Recognition of group membership by voice in Mexican jays, Aphelocoma ultramarina. Animal Behaviour, 62(2), 297–303. 10.1006/anbe.2001.1745

Jiguet, F. (2020). The Fox and the Crow. A need to update pest control strategies. Biological Conservation, 248(July), 108693. 10.1016/j.biocon.2020.108693

Kaplan, G. (1999). Song structure and function of mimicry in the Australian magpie (Gymnorhina tibicen): Compared to lyrebird (Menura ssp.) International Journal of Comparative Psychology.

Keenan, S., Mathevon, N., Stevens, J. M., Nicolè, F., Zuberbühler, K., Guéry, J. P., & Levréro, F. (2020). The reliability of individual vocal signature varies across the bonobo’s graded repertoire. Animal Behaviour, 169, 9–21. 10.1016/j.anbehav.2020.08.024

Kelley, J. L., Morrell, L. J., Inskip, C., Krause, J., & Croft, D. P. (2011). Predation risk shapes social networks in fission-fusion populations. PLoS ONE, 6(8). 10.1371/journal.pone.0024280

Kershenbaum, A., Blumstein, D. T., Roch, M. A., Akçay, Ç., Backus, G., Bee, M. A., Bohn, K., Cao, Y., Carter, G., Cäsar, C., Coen, M., Deruiter, S. L., Doyle, L., Edelman, S., Ferreri-Cancho, R., Freeberg, T. M., Garland, E. C., Gustison, M., Harley, H. E., … Zamora-Gutierrez, V. (2014). Acoustic sequences in non-human animals: A tutorial review and prospectus. Biological Reviews, 91(1), 000–000. 10.1111/brv.12160

Kondo, N., Izawa, E. I., & Watanabe, S. (2010). Perceptual mechanism for vocal individual recognition in jungle crows (Corvus macrorhynchos): Contact call signature and discrimination. Behaviour, 147(8), 1051–1072. 10.1163/000579510X505427

Kondo, N., & Watanabe, S. (2009). Contact calls: Information and social function. Japanese Psychological Research, 51(3), 197–208. 10.1111/j.1468-5884.2009.00399.x

Konishi, M. (1985). Birdsong: From behavior to neuron. 10.1146/annurev.neuro.8.1.125

Kremers, D., Lemasson, A., Almunia, J., & Wanker, R. (2012). Vocal sharing and individual acoustic distinctiveness within a group of captive orcas (Orcinus orca). Journal of Comparative Psychology, 126(4), 433–445. 10.1037/a0028858

Laiolo, P., Palestrini, C., & Rolando, A. (2000). A study of Choughs’ vocal repertoire: Variability related to individuals, sexes and ages. Journal fur Ornithologie, 141(2), 168–179. 10.1046/j.1439-0361.2000.00074.x

Lenth, R. V. (2023). Emmeans: Estimated marginal means, aka least-squares means [R package version 1.8.8]. https://CRAN.R-project.org/package=emmeans

Linhart, P., Osiejuk, T. S., Budka, M., Šálek, M., Špinka, M., Policht, R., Syrová, M., & Blumstein, D. T. (2019). Measuring individual identity information in animal signals: Overview and performance of available identity metrics. Methods in Ecology and Evolution, 10(9), 1558–1570. 10.1111/2041-210X.13238

Marler, P. (2004). Bird calls: a cornucopia for communication. In Nature’s music (pp. 132–177). Elsevier.

Marler, P., & Tamura, M. (1962). Song “dialects” in three populations of white-crowned sparrows. The Condor, 64, 368–377.

Marshall, A. J., & Coombs, C. J. (1957). The Interaction of Environmental, Internal and Behavioural Factors in the Rook, Corvus F. Frugilegus Linnaeus. Proceedings of the Zoological Society of London, 128(4), 545–588. 10.1111/j.1096-3642.1957.tb00275.x

Martin, K., Cornero, F. M., Clayton, N. S., Adam, O., Obin, N., & Dufour, V. (2024). Vocal complexity in a socially complex corvid : gradation, diversity and lack of common call repertoire in male rooks. Royal Society Open Science, 11. 10.1098/rsos.231713

Mates, E. A., Tarter, R. R., Ha, J. C., Clark, A. B., & McGowan, K. J. (2015). Acoustic profiling in a complexly social species, the American crow: Caws encode information on caller sex, identity and behavioural context. Bioacoustics, 24(1), 63–80. 10.1080/09524622.2014.933446

McGregor, P. K. (1980). Song Dialects in the Corn Bunting (Emberiza calandra). Zeitschrift für Tierpsychologie, 54(3), 285–297. 10.1111/j.1439-0310.1980.tb01246.x

McInnes, L., Healy, J., & Melville, J. (2018). UMAP: Uniform manifold approximation and projection for dimension reduction. arXiv.

Monteiro, R. d. A., Ferreira, C. D., & Perbiche-Neves, G. (2021). Vocal repertoire and group-specific signature in the smooth-billed ani, crotophaga ani linnaeus, 1758 (Cuculiformes, aves). Papeis Avulsos de Zoologia, 61, 1–9. 10.11606/1807-0205/2021.61.59

Mundinger, P. C. (1982). Microgeographic and macrogeographic variation in the acquired vocalizations of birds. Acoustic communication in birds.

Patterson, I. J., Dunnet, G. M., & Fordham, R. A. (1971). Ecological Studies of the Rook, Corvus frugilegus L., in North-East Scotland: Dispersion. The Journal of Applied Ecology, 8(3), 815. 10.2307/2402685

Peckre, L., Kappeler, P. M., & Fichtel, C. (2019). Clarifying and expanding the social complexity hypothesis for communicative complexity. Behavioral Ecology and Sociobiology, 73(1). 10.1007/s00265-018-2605-4

Průchová, A., Jaška, P., & Linhart, P. (2017). Cues to individual identity in songs of songbirds: testing general song characteristics in Chiffchaffs Phylloscopus collybita. Journal of Ornithology, 158(4), 911–924. 10.1007/s10336-017-1455-6

R Core Team. (2021). R: A Language and Environment for Statistical Computing. https://www.r-project.org/

Richardson, S. C., Patterson, I. J., & Dunnet, G. M. (1979). Fluctuations in Colony Size in the Rook, Corvus frugilegus. Journal of Animal Ecology, 48(1), 103–110. http://www.jstor.org/stable/4103

Röell, A., & Bossema, I. (1982). A comparison of nest defence by Jackdaws, rooks, magpies and crows. Behavioral Ecology and Sociobiology, 11(1), 1–6. 10.1007/BF00297658

Roskaft, E., & Espmark, Y. (1982). Vocal communication by the rook ¡i¿Corvus frugilegus¡/i¿ during the breeding season. Ornis Scandinavica, 13(1), 38–46. 10.2307/3675971

Røskaft, E., & Espmark, Y. (1984). Sibling recognition in the rook (¡i¿Corvus frugilegus¡/i¿). Behavioural Processes, 9(2-3), 223–230. 10.1016/0376-6357(84)90042-1

Sainburg, T., & Gentner, T. Q. (2021). Toward a Computational Neuroethology of Vocal Communication : From Bioacoustics to Neurophysiology, Emerging Tools and Future Directions. 15(December), 1–24. 10.3389/fnbeh.2021.811737

Salinas-Melgoza, A., & Wright, T. F. (2012). Evidence for vocal learning and limited dispersal as dual mechanisms for dialect maintenance in a parrot. PloS one, 7(11). 10.1371/journal.pone.0048667

Searcy, W. A. (1992). Song repertoire and mate choice in birds. Integrative and Comparative Biology, 32(1), 71–80. 10.1093/icb/32.1.71

Sewall, K. B. (2009). Limited adult vocal learning maintains call dialects but permits pair-distinctive calls in red crossbills. Animal Behaviour, 77(5), 1303–1311. 10.1016/j.anbehav.2009.01.033

Slabbekoorn, H., & Peet, M. (2003). Birds sing at higher pitch in urban noise. Nature, 424(July), 267.

Slater, P. J., & Ince, S. A. (1982). Song Development in Chffinches: What Is Learnt and When? Ibis, 124(1), 21–26. 10.1111/j.1474-919X.1982.tb03737.x

Smith-Vidaurre, G., Araya-Salas, M., & Wright, T. F. (2020). Individual signatures outweigh social group identity in contact calls of a communally nesting parrot. Behavioral Ecology, 31(2), 448–458. 10.1093/BEHECO/ARZ202

Stowell, D., Morfi, V., & Gill, L. F. (2016). Individual identity in songbirds: Signal representations and metric learning for locating the information in complex corvid calls. Proceedings of the Annual Conference of the International Speech Communication Association, INTERSPEECH, 08-12-Sept, 2607–2611. 10.21437/Interspeech.2016-465

Swingland, I. R. (1977). The social and spatial organization of winter communal roosting in Rooks (Corvus fmgilegus). Journal of Zoology, 182(4), 509–528. 10.1111/j.1469-7998.1977.tb04167.x

Terry, A. M., Peake, T. M., & McGregor, P. K. (2005). The role of vocal individuality in conservation. Frontiers in Zoology, 2, 1–16. 10.1186/1742-9994-2-10

Thomas, M., Jensen, F. H., Averly, B., Demartsev, V., Manser, M. B., Sainburg, T., Roch, M. A., & Strandburg-Peshkin, A. (2022). A practical guide for generating unsupervised, spectrogram-based latent space representations of animal vocalizations. Journal of Animal Ecology, 91(8), 1567–1581. 10.1111/1365-2656.13754

Vignal, C., Mathevon, N., & Mottin, S. (2008). Mate recognition by female zebra finch : Analysis of individuality in male call and first investigations on female decoding process. Behavioural Processes, 77, 191–198. 10.1016/j.beproc.2007.09.003

Wascher, C. A., Szipl, G., Boeckle, M., & Wilkinson, A. (2012). You sound familiar: Carrion crows can differentiate between the calls of known and unknown heterospecifics. Animal Cognition, 15(5), 1015–1019. 10.1007/s10071-012-0508-8

Willis, C. K., & Brigham, R. M. (2004). Roost switching, roost sharing and social cohesion: Forest-dwelling big brown bats, Eptesicus fuscus, conform to the fission-fusion model. Animal Behaviour, 68(3), 495–505. 10.1016/j.anbehav.2003.08.028

Wright, T. F., & Dahlin, C. R. (2017). Vocal dialects in parrots: Patterns and processes of cultural evolution. Emu, 118(1), 50–66. 10.1080/01584197.2017.1379356

Yorzinski, J. L., Vehrencamp, S. L., McGowan, K. J., & Clark, A. B. (2006). The Inflected Alarm Caw of the American Crow: Differences in Acoustic Structure Among Individuals and Sexes. The Condor, 108(3), 518–529.

