## Supplementary material for "Can we trace the social affiliation of rooks (*Corvus frugilegus*) through their vocal signature?": Cartography of wild colonies with sampled nests

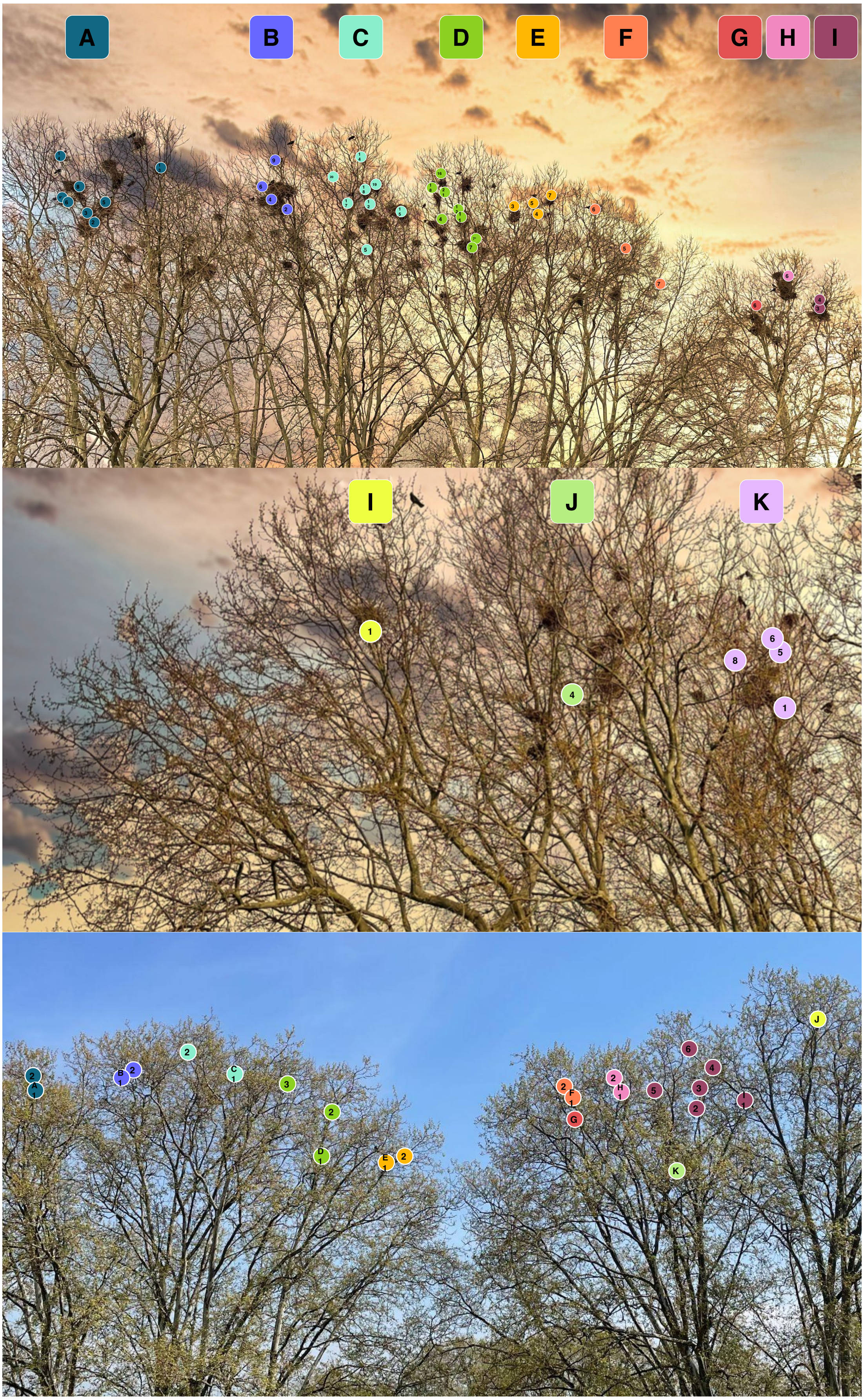


Figure S1. Cartography of the sampled nests in the 'Strasbourg wild' (top and middle, photo split due to the size of the colony) and 'Contades' (bottom) colonies, showing the individual nests and the nest clusters defined in the article. Each colour corresponds to a nest cluster, with nests numbered in the order they were sampled, separately for each nest cluster.
